# Active Fixation as an Efficient Coding Strategy for Neuromorphic Vision

**DOI:** 10.1101/2022.09.29.510091

**Authors:** Simone Testa, Silvio P. Sabatini, Andrea Canessa

## Abstract

Contrary to a photographer, who puts a great effort in keeping the lens still, eyes insistently move even during fixation. This benefits signal decorrelation, which underlies an efficient encoding of visual information. Yet, camera motion is not sufficient alone; it must be coupled with a sensor specifically selective to temporal changes. Indeed, motion induced on standard imagers only results in burring effects. Neuromorphic sensors represent a valuable solution. Here we characterize the response of an event-based camera equipped with *Fixational Eye Movements* (FEMs) on both synthetic and natural images. Our analyses prove that the system starts an early stage of redundancy suppression, as a precursor of subsequent whitening processes on the amplitude spectrum. This does not come at the price of corrupting structural information contained in local spatial phase across oriented axes. Isotropy of FEMs ensures proper representations of image features without introducing biases towards specific contrast orientations.

## Introduction

Human vision rapidly adapts to unchanging retinal input up to experiencing a real perceptual fading when retinal image motion is artificially compensated or eliminated. Also for this reason, vision is an active process and this need leads our eyes to constantly move for keeping motionless parts of the visual scene visible. A particular class of involuntary eye movements, known as Fixational Eye Movements (FEMs), serves this purpose^1^.

In the attempt of understanding how brain exploits eye’s jitter, neuromorphic engineering and event-based vision sensors^2,3^ provide a natural “learning-by-doing” framework to investigate the early stages of visual processing in active (i.e., real world) conditions^4^. These cameras convert a visual scene into a stream of asynchronous ON and OFF events based on positive or negative temporal contrast; as opposed to frame-based and clock-driven acquisitions of luminance. These continuous-time sensors functionally emulate the key features of the human retina and represent a major shift from conventional cameras, by transmitting only pixel-level changes at microsecond precision. It is therefore not surprising that, as for neurons in the retina, no visual information can be gained in the absence of relative motion between the sensor and the environment. An active vision mechanism based on FEMs can be implemented on a bio-inspired robotic system for making visual perception of static objects feasible by event-based sensors.

Besides being a means for refreshing neural activity and preventing perceptual fading (retinal adaptation), FEMs have been pinpointed to play a key role in terms of efficiency coding^5^. The efficiency principle^6^ states that one of the goals of early vision processing is to maximize the information that is encoded about relevant sensory variables, given constraints on the available (neural) resources (e.g., the limited capacity of the optic nerve), by reducing uninformative correlations typical of natural scenes. Before advancing this hypothesis, the spatio-temporal behavior of retinal bipolar and ganglion cells (RGCs) has long been considered as the only responsible for this signal decorrelation. At first approximation, RGCs act as linear spatio-temporal filters that implement lateral and temporal inhibition to generate receptive fields with antagonist center-surround spatial organization and transient (i.e., biphasic) response in time. In this way, they seek to reduce redundancy between parallel channels in space, and within each single channel along time. In addition to that, contributions of non-linear stimulus-response relationships^7^ (such as synaptic rectification, depression, gain control, spiking threshold and refractory) refine the job, eventually permitting retinal neurons to transmit information with nearly optimal efficiency. However, this view lacks to consider the observer’s motor activity^8^, relying on the simplifying assumption that the input to the retina is a stationary image, or - at the most - a sequence of stationary frames. In living animals, the retina receives unstable visual inputs caused by movements of body, head and eyes. Even when an animal is fixating an object, the whole image on the retina is shifted by the presence of incessant microscopic albeit continuous and erratic eye movements. In Segal *et al*.^8^, authors proved that the response of RGCs alone still exhibits strong and extensive spatial correlations in the absence of fixational eye movements (e.g., with stimulus flash). In the presence of FEMs, instead, the levels of correlation in the neural responses dropped significantly, resulting in effective decorrelation of the channels streaming information to the brain. These observations demonstrate that microscopic eye movements act to reduce correlations in retinal responses and contribute to visual information processing. Similar conclusions have also been drawn in^9^. They demonstrated that the statistics of FEMs matches the statistics of natural images, such that their interaction generates spatiotemporal inputs optimized for processing by RGCs. This spatiotemporal reformatting is crucial for neural coding, as it matches the range of peak spatiotemporal sensitivity of retinal neurons in primates. As a consequence, jittery movements of a sensor can emphasize edges, as postulated and formalized in the fascinating theory of the *Resonant Retina*^10^, and very recently further examined in^11^.

In the present research, we investigate the effects of fixational eye movements on neuromorphic sensors. Given the low occurrence and supposed minor significance of micro-saccades^9^ as compared to the major effects due to slow fixational drifts (which are the two main components of FEMs), we target non-saccadic FEMs only. We extend an existing model of biological fixational movements^12^ to increase their isotropy. We use this model to move a neuromorphic camera in a biological fashion while acquiring data from synthetic and natural stimuli. The resulting event stream is analyzed for characterizing the role of FEMs. The main focus is on understanding how it preserves structural information of the input natural images while decorrelating their amplitude in order to reduce redundancy.

## Results

### FEM simulations

Inducing bio-inspired fixational eye movements on an artificial sensor requires, first of all, a good model of such movements based on their known characteristics in natural viewing. Although Brownian motion, with its erratic trajectory, is frequently assumed as a valid model for approximating FEM, more accurate mathematical models can capture other fundamental properties of fixational instability^13^. We have therefore chosen the *Self-Avoiding random Walk* (SAW)^12^ for simulating our FEM paths. However, we adapted such model in order not to limit each step towards the cardinal directions, but admitting a wider set of possible orientations. This leads towards an isotropic and more biologically plausible visual exploration around the fixation point. An example of the resulting FEM sequence generated in this way is shown in Figure 1A (as well as in the right panels of Figure 4). The same figure (see panel C) displays how the DVS activity roughly reflects the FEM sequence followed by the sensor during acquisition: the firing activity is phase locked to the movement and its amplitude relates to that of the underlying FEM steps.

**Figure 1.**
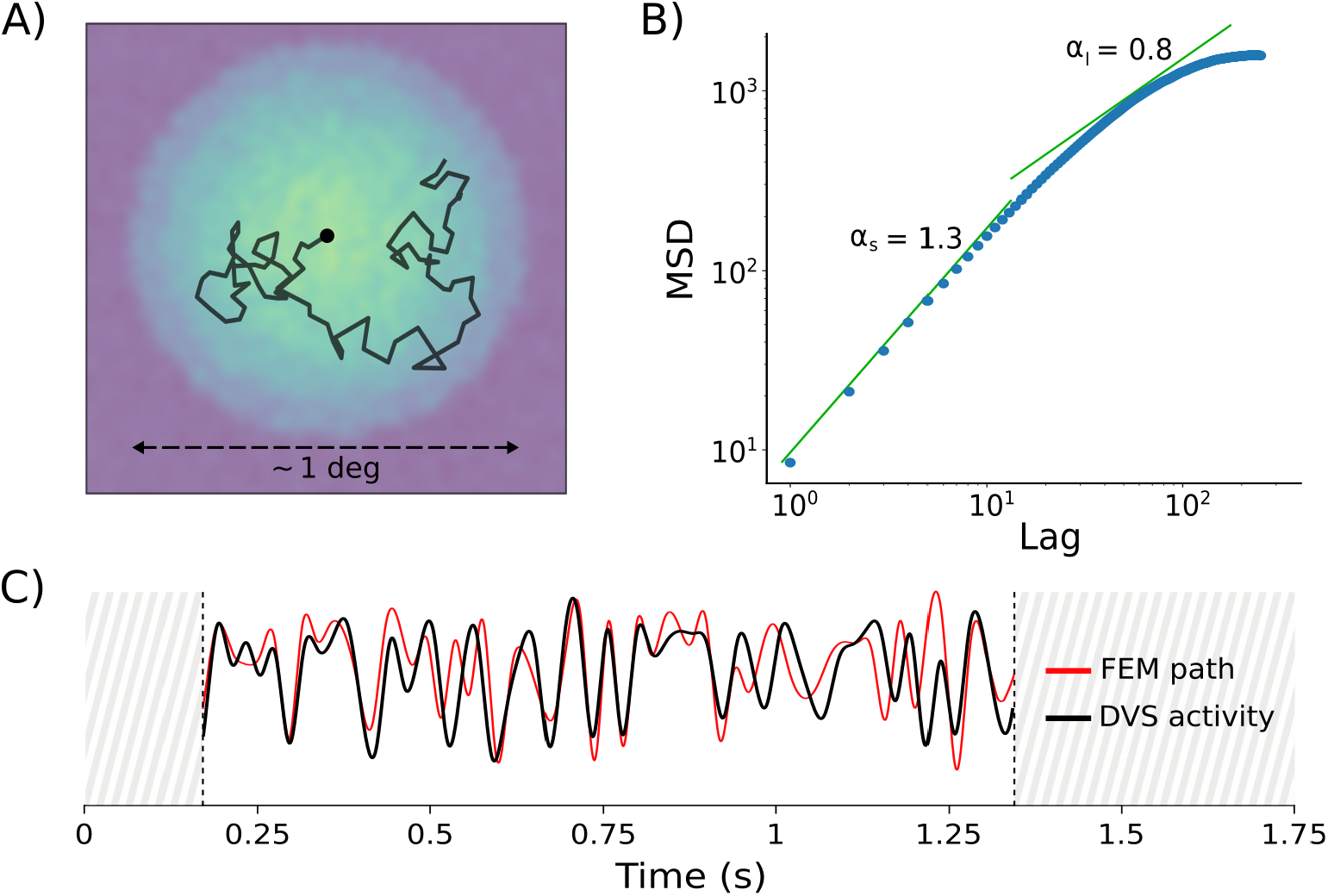
Characterization of the proposed model, adapted from^12^. A) FEM sequence obtained by the model. Black line represents an example of FEM trajectory with 80 steps. The FEM path is superimposed on its activation field, where the greenish shades of blue indicate lower activation values: the circular shape reveals the region of the foveola in which FEMs are confined. B) Temporal evolution of the mean squared (spatial) displacement (in arcmin^2^) as a function of the iteration (or time) lag. C) The distance covered by the walker during a specific FEM sequence (specific seed) is shown in red (interpolation over 60 FEM steps). The (smoothed) instantaneous firing rate of the DVS (averaged across 16 different recordings) is superimposed in black, sampled at all 60 steps and interpolated. Signals are shown only for the time window of fixational eye movements.

Since we modified the original model, we tested whether some of its major predictions were still maintained. Specifically, the original model replicated both persistence and anti-persistence behaviors on short and long timescales respectively, well matching some experimental evidence from biology^14^. In biological data, the mean squared displacement (MSD) has a power-law trend with the lag *l*: persistence is exhibited on a short timescale with a fitting exponent *α*_*s*_ *>* 1 (∼1.4) and anti-persistence on a long timescale with *α*_*l*_ *<* 1 (∼0.8)^14^. In spite of the changes we made, we still observe similar results (cf.^12^) with our modified version of the model (1B): persistence is visible on a short timescale since the power-law exponent of the mean squared displacement with respect to iteration lags is *α*_*s*_ = 1.3 *>* 1 (lag ≤10 iterations), while *α*_*l*_ = 0.8 *<* 1 denotes anti-persistence on the long timescale (lag *>* 10 iterations).

### Decorrelation of natural images

In order to study whitening effects of FEMs on our neuromorphic platform, we first of all reproduced similar experiments as^9^, but analyzing the output of the system instead than the power of the dynamic retinal input. We expect that the activity elicited in each receptor of our artificial retina complies with the statistical properties of natural images in the same way it happens in biology. In case of stimuli with a fixed contrast value, results showed that pixels’ mean activity grows as spatial frequency increases (see^15^ and Supplementary Fig.S1 online). As a matter of fact, by randomly moving around, each receptor scans an increasing number of edges as the spatial frequency grows, thus eliciting an increasing number of events. Remarkably, by adjusting gratings’ contrast according to the 1*/k* falloff of natural image amplitude spectrum (with *k* representing spatial frequency), the response of the system gave a roughly constant firing rate over the whole range of frequency. This whitening effect is attributable to the opposing trend between the amplitude distribution across spatial frequencies (typical of natural images) with respect to the amplification introduced by FEMs.

We then compared the decorrelation effect of the neuromorphic active-vision system to classical (frame-based) whitening techniques on images from the natural world. The traditional methods we considered were (i) the whitening approach proposed by Olshausen and Field^16^ (which we will refer to as OFW from now on) and (ii) the *Difference of Gaussians* (DOG) filtering technique - details of these decorrelation strategies are given in Methods section. Concerning event-based recordings, instead, we can distinguish them based on stimulation procedure: (i) flash-based (static camera, flashing stimulus) and (ii) FEM-based (static stimulus, shaking camera). In order to compare such recordings to frame-based whitened images, event pre-processing was required for building frames by integration in a proper time window, see Methods for details. Interestingly, the optimal value of this time window (∼ 200 ms) matches with the average duration of fixation as reported in the literature^17^, suggesting a possible relation. Examples of the resulting images are shown in Figure 2A, together with the original natural image (from van Hateren dataset^18^) for comparison. From a first qualitative (visual) inspection of these different decorrelation strategies, we notice remarkable similarities between the natural image whitened according to^16^ and that obtained by filtering it with a DOG kernel (ON-center), as expected. However, less intuitively, the image reconstructed from DAVIS events recorded under FEMs shows some similarity as well. With a flashing stimulus, instead, the resulting image more faithfully resembles the original image, suggesting that strong and uninformative correlations between signals carried by different pixels - typical of natural images - still remain.

**Figure 2.**
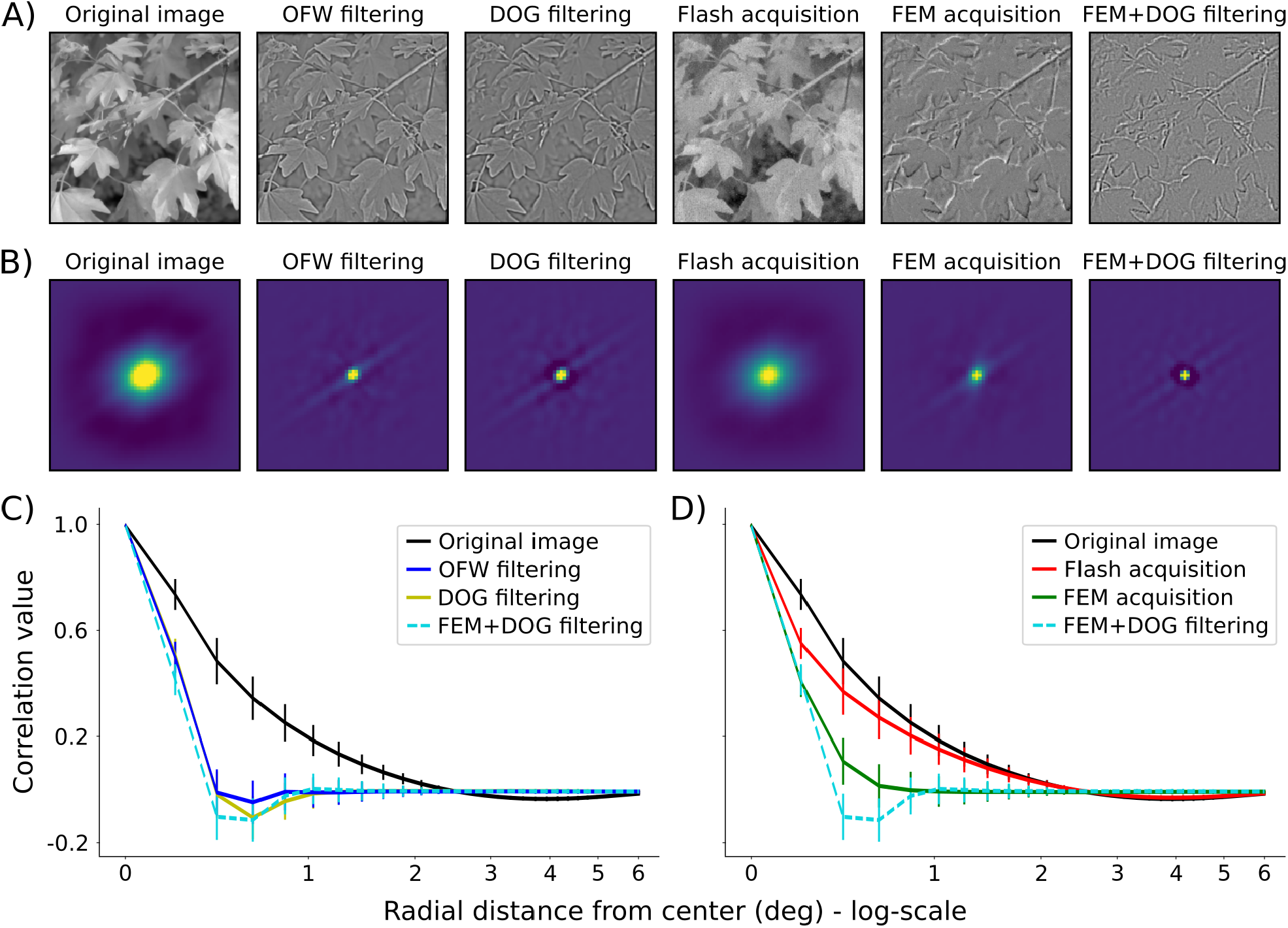
Amplitude information analysis. A) Effects of different “frame-based” whitening procedures compared to “event-based” counterparts, and their combination. A data sample from the van Hateren dataset (cropped and down-sampled to 200 ×200) is shown in the left-most image. The two consecutive frames represent results from standard whitening techniques, while the next two images are reconstructed from event-based recordings by either flashing the stimulus on the monitor or shaking the camera with FEM-based sequences. The cascade of a DOG band-pass filtering on the FEM-based reconstructed frame is shown in the right-most image. All axes range from −20 deg to 20 deg. B) Two-dimensional auto-correlations of the images in panel A with matching positions. Axes range from −6 deg to 6 deg. C) and D) Comparison of the azimuthal-averaged profiles of the 2D auto-correlations. Each curve represents mean and standard deviation across the whole set of natural images, also averaged across FEM seeds (or flash trials) in case of event-based acquisitions. For the sake of clearness, we split in C) the results from standard whitening techniques (blue and yellow lines) while in D) the results from reconstructed images of both event-based acquisitions (red and green lines). The average correlation profile of the original image (black line) and the combination of FEM effects with DOG filtering (dashed light-blue line) are displayed in both panels for comparison.

A quantitative measure of the decorrelation effect is revealed from a second-order statistical analysis: specifically, by estimating the auto-correlation functions of the corresponding images (see Figure 2B). We can observe that data obtained by FEMs seem to be as highly decorrelated as standard techniques, while correlation values of flash-based recordings are very close to those of the original image. These results can be better appreciated by looking at the azimuthal average of such 2D correlation functions, as shown in Figure 2C and D. Here, we can notice that the auto-correlation function is more sharply peaked in zero lag both (i) when standard whitening procedures are applied on the original image (blue and yellow curves in panel C) or (ii) when the image is captured by a shaking neuromorphic camera (green curve in panel D). Conversely, in case of image flashes recorded by the same sensor, a high correlation value is shown also from pairs of pixels far apart one another (red), similarly to the original image (black). In other words, the shaking neuromorphic sensor, as well as standard whitening procedures, highly decorrelates the input signal, only preserving non-redundant information. Image flash, instead, preserves redundancies. As a matter of fact, flashing the stimulus on monitor and recording it with a static sensor is somehow equivalent to a rate-based encoding of the image: each pixel outputs a spike train with firing rate proportional to the gray-scale value in the corresponding location of the original image. Finally, by applying a DOG filtering in cascade to the FEM-based acquisition, we can notice that fixational movements boost the decorrelation induced by RGC-like filters (light blue curve in both panels).

### Preservation of phase

The auto-correlation function alone is not exhaustive for defining the system as an efficient encoder since it relates to amplitude information, only. In this case, nothing can be inferred on whether the structure of the image is preserved, which is revealed by phase information^19^. Specifically, while redundant amplitude information must be neglected in an efficient coding framework, phase information must be preserved since it encodes the characteristic image structures, as spatial symmetries of contrast. Hence, it is worthwhile examining phase properties both at a global and local scale. A very coarse metric for evaluating structure-related information is given by the phase correlation coefficient, i.e. the correlation of global phase between a reference image and its filtered version. We used this metric to find out which one between the two classical whitening procedures should be considered as the gold standard for the preservation of phase information, in order to take it as a reference for further and more accurate analysis. We found out that global phase is totally preserved in the OFW technique proposed by^16^ since the correlation coefficient between the phase spectrum of the original image and that of the OFW-based image is very close to unit (*>* 0.99). Instead, global phase is not completely preserved in the DOG-based decorrelation strategy since correlation coefficient equals to 0.8. Therefore, all the results from subsequent analysis on local phase in event-based recordings will be referred to phase information in the OFW-filtered image. A bank of Gabor filters was then applied on both the reconstructed FEM- and flash-based event images in order to compare the dominant local phase (details in the phase-based analysis paragraph of the Methods section). Results are shown in Figure 3A for a Gabor filter with peak frequency of ∼ 0.5 cyc/deg.

**Figure 3.**
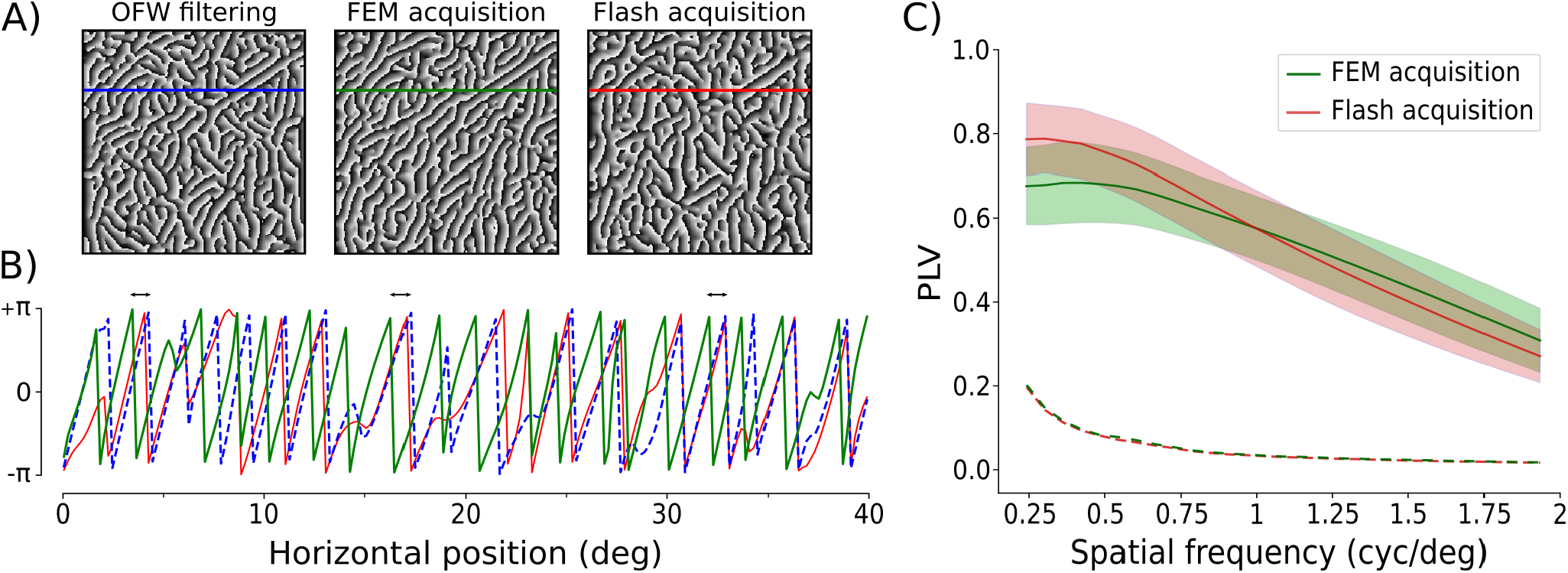
Phase information analysis. A) Results of the dominant local phase extracted at ∼0.5 cyc/deg from three images in Figure 2A. Axes range from −20 deg to 20 deg. B) Mono-dimensional view of the dominant local phase extracted at a specific row. Colors of the curves match those of horizontal lines superimposed on the top three images: phase from FEM (green) and flash (red) acquisitions must be compared to that from the reference whitening procedure (dashed blue). Double-head arrows on top of the curves point out some examples of the roughly-constant phase shift between the reference and FEM-based image, yet not affecting the conservation of phase structure. C) Distribution of the PLV for different spatial frequencies. The mean PLV (with respect to the OFW-filtered image) are shown with green and red lines for the FEM-based and flash-based signals respectively. Standard deviations are visible as shaded areas. Dashed curves represent the statistical significance level (95-percentile confidence interval) obtained by the PLV distribution of the surrogates for both FEM- (green) and flash-based (red) images.

In order to quantify the consistency of the detected dominant local phase in event-based signals with respect to the reference whitened frame, we compute their *Phase Locking Value* (PLV)^20^. This metrics allows to assess the preservation of phase structure between the reference and the event-reconstructed image despite some possible phase shift between the two signals, which could be relevant in FEM-based recordings due to the motion of the sensor (as highlighted in Figure 3B). In Figure 3C we compare the PLV (averaged across all sets of recordings) of FEM- and flash-based data at different scales of the Gabor filters (from ∼ 0.2 to 2 cyc/deg, as for the cutoff frequency of the OFW method). The PLVs of the reconstructed images from FEM-based recordings are high (in a [0, 1] range) for a broad range of tested spatial frequencies, proving the preservation of the underlying image structure. Similar results are observed, as expected, in case of flash-based recordings, which keep all spatial information of the original image. It is worth noting that most of the energy in natural images is confined at lower frequencies - approximately up to ∼ 0.5 cyc/deg - as it results from their power spectra. Therefore, it is not surprising that, after such value, the PLV decays more sharply. Furthermore, the PLV curve of FEM-based images is flatter than its flash-based counterpart, suggesting an increased reliability of phase information across a wider range of spatial frequencies as a consequence of whitening^21^. Finally, all such values are statistically significant against a permutation test based on surrogate data for all frequencies.

### Importance of isotropy

The last experiment we made to prove possible benefits of FEMs in neuromorphic vision relates to the isotropic character of such peculiar motion sequence. First of all, SAW model resulted in a population of trajectories characterized by angular directions uniformly distributed around the circle (*P* = .42, Rayleigh test), while for all biased movements we reject the null hypothesis (*P <* .001, Rayleigh test). Similar results were achieved from a circular uniformity test (biased movements: *P <* .001 from the *vtest()* function in *pycircstat*.*tests*; SAW-based movements: *P* = .15 from *vtest()*). Finally, a symmetry test proved that the unbiased movement is symmetric about the median (*P* = .72 from *symtest()* function), while this is not true for biased movements (*P <* .001 from *symtest()*). We can therefore conclude that the SAW-based motion sequences could be considered as isotropic, not showing any remarkable bias. It is worth noting that the longer is the duration of active fixations (i.e. the more FEM steps are performed), the greater is the capability of detecting all oriented contrasts, in line with what has been observed in visual acuity experiments^22^.

We then analyzed the response of the sensor to images of differently oriented gratings either in the presence of isotropic or anisotropic movements. Examples of the different FEM paths are shown in Figure 4 for different orientation biases. Results prove the effectiveness of FEMs’ isotropy in equalizing sensor’s response with respect to all possible orientations of visual stimuli. We observed that, for direction-biased movements, the response of the system was different at various orientations of the stimulus - as visible from the blue curves in all three plots of Figure 4. Specifically, the maximum firing rate was achieved when the grating had an angle shifted by 90° with respect to the direction bias of the motion, meaning that a bias in the FEM sequence reflects in a preferred orientation detection. On the contrary, the black curve in the same plots shows that the neuromorphic camera exhibited equalized activity with respect to differently oriented gratings when scanning all possible directions with isotropic erratic movements.

**Figure 4.**
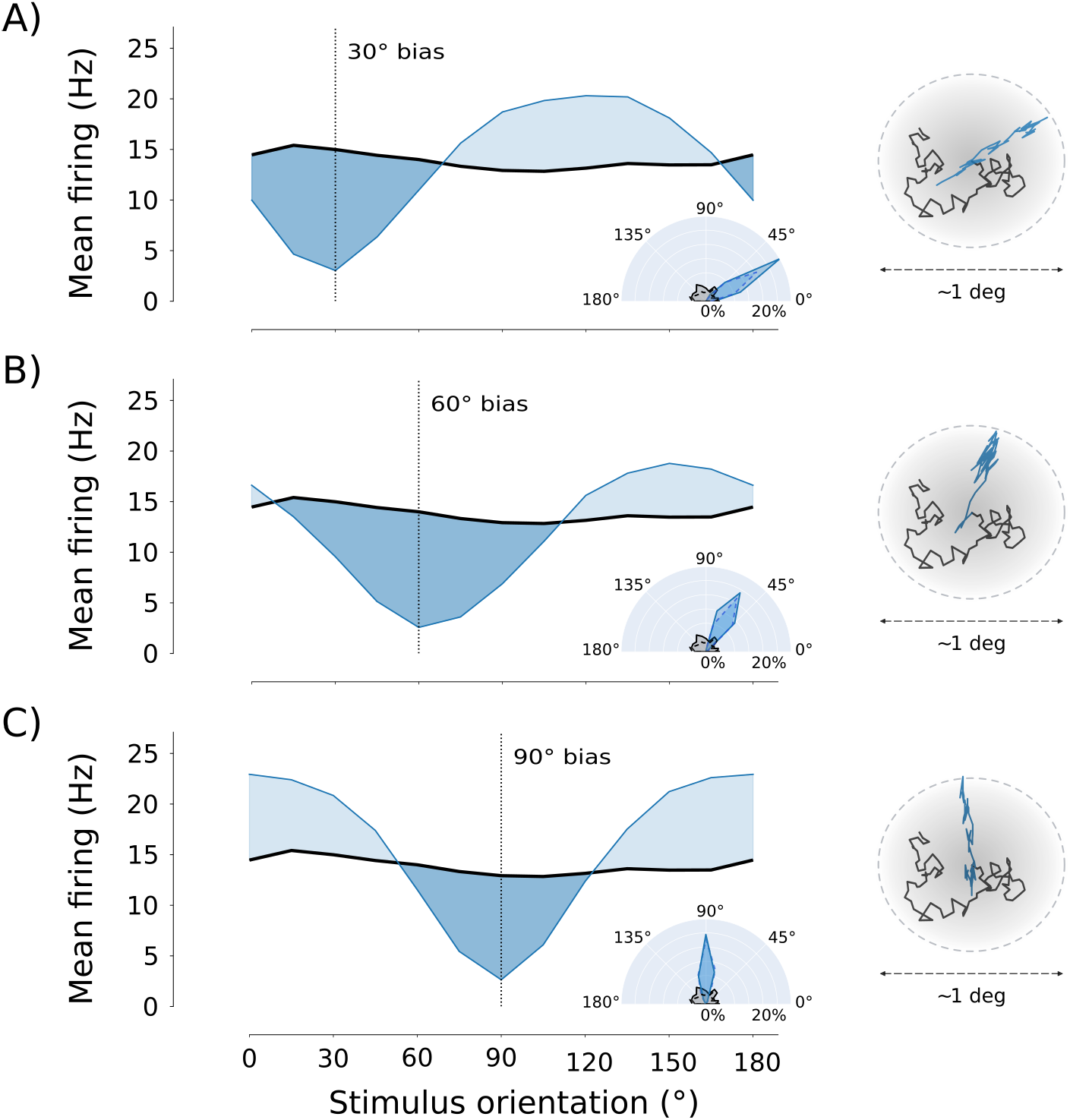
Effect of motion (an)isotropy on DVS response. A) Comparison of the mean firing rate over the sensor area evoked by 30° biased motion sequences (blue) and SAW-based isotropic (black) FEMs. The right insets represent examples of anisotropic (blue) and isotropic (black) trajectories on the top, and their corresponding circular histograms on the bottom. Panels B) and C) are same as A) but for 60° and 90° bias, respectively.

### FEM steps as a spatial filtering stage

The event-based sensor, as the retinal neurons by which it has been inspired, is mainly sensitive to time derivative of luminance. In the presence of a relative motion between the sensor and the scene, such derivative relates to spatial gradients in the image (see equation 1). One main property of image gradient is its local orientation. In case of a static scene, such an orientation is mostly represented when it is perpendicular to the sensor’s direction of movement. By splitting a FEM pattern in all the single movements (steps) that compose it, we can distinguish their individual contribution to the representation of visual information. As a matter of fact, each frame is the result of the operation produced by the neuromorphic sensor when subject to an oriented movement. Single FEM steps on the silicon retina act as anisotropic oriented band-pass filters applied on the underlying image. This can be seen in Figure 5A that shows a set of 36 examples of such filters in Fourier domain. All filters were reconstructed by solving a Tikhonov minimization problem from an equivalent number of steps, related to the same FEM seed. When taken together, the combination of all differently-oriented kernels builds up an isotropic filter (see Figure 5B), which is comparable to a DOG profile in the frequency domain.

**Figure 5.**
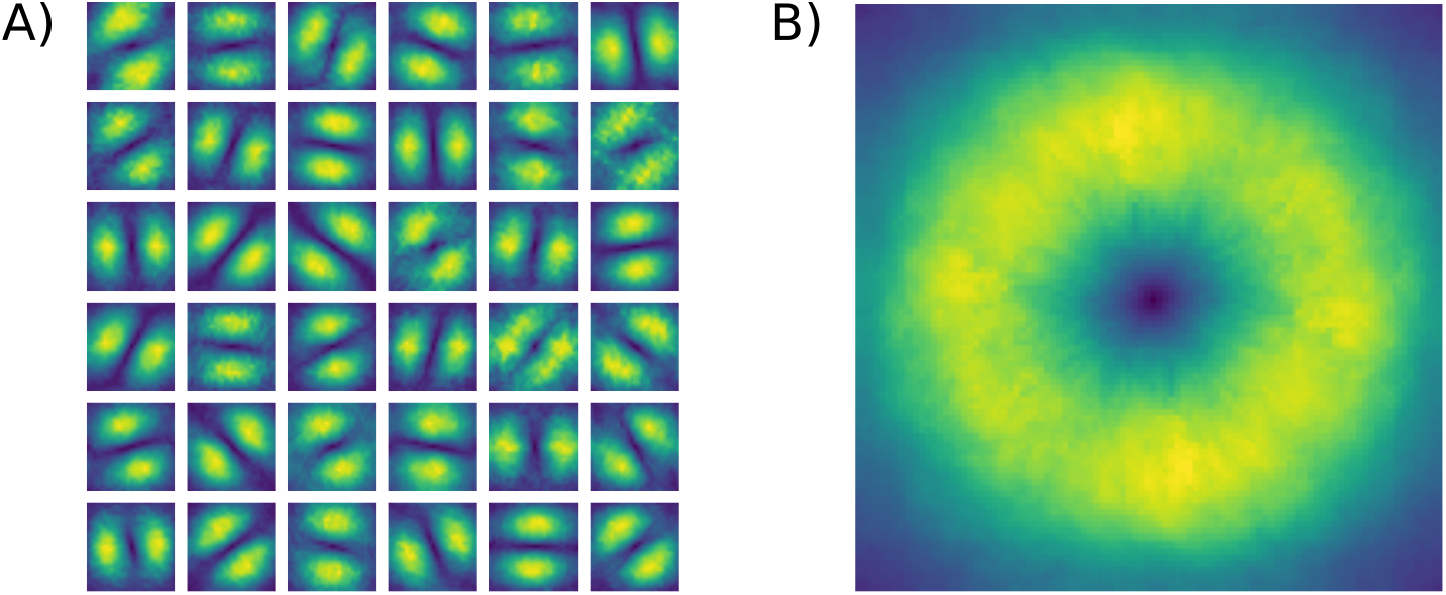
Equivalent FEM spatial filters. A) Examples of the anisotropic filters in Fourier domain achieved from single FEM steps at a given seed and averaged across all natural image stimuli. B) Overall isotropic filter obtained by averaging across the whole FEM sequence.

## Discussion and conclusions

Despite the name, human fixation is a highly dynamic process. In biology, some roles of fixational eye movements have already been pointed out and discussed, persuading the scientific community that FEMs are far from being a nuisance, as originally believed. In this work, we investigate the role of FEMs in neuromorphic vision, i.e on the output signal of silicon circuits that emulate some primary functionalities of the human retina (namely, transient dynamics, no spatial filtering).

We started our investigation by examining the overall spectral response of the system following a similar procedure as in^9^. While the power spectrum in natural scenes is highly concentrated at low spatial frequencies (with an amplitude falloff of 1*/k*), a neuromorphic and actively-fixating system intrinsically enhances higher spatial-frequency contents by amplifying its response to them. Therefore, such a system tends to oppose to the power-law falloff, counterbalancing the latter and enabling an equalized response to all discernible frequencies when the stimulus has such statistical properties. The investigation with natural image stimuli also proved that our neuromorphic system equipped with FEMs starts an early stage of redundancy suppression as a precursor of subsequent whitening processes^23–25^. Since no explicit spatial filtering is actually implemented from the DAVIS sensor, the origin of the observed whitening effect should be ascribed to the combination of three main characteristics of the sensing strategy: (1) the peculiar motion used^8^, (2) the transient response of the camera, and (3) some non-linear behavior^7^ in the acquisition process of single pixels. However, when the neuromorphic camera recorded the same natural image flashing on the monitor, the resulting signal was still highly correlated despite intrinsic non-linearity of the sensor and trial-to-trial noise in the recordings. Therefore, much of the decorrelation in the FEM-based signal is ascribable to the combination of movements and sensitivity to brightness transitions, with a small contribution of additive noise and non-linear behaviors. In other words, the small image displacements induced by FEMs - given retina/sensor temporal DC removal - help discarding redundant spatial correlations of natural visual input, hence boosting the decorrelation induced by subsequent center-surround filtering of RGCs (not sufficient alone to disrupt the strong correlations of natural images^7^).

A mere weak correlation of the input signal does not yet imply a highly efficient coding system. For instance, if two signals are affected by independent noise, this decorrelates them without improving coding efficiency. In order to efficiently encode a visual scene, it is necessary that the decorrelation procedure does not compromise the preservation of its structure-related information. Despite being commonly related to coding efficiency, second-order statistical moments such ad the autocorrelation function and its Fourier counterpart - the power spectrum - consider only amplitude information, by definition insensitive to the local phase, which conveys most of the information about image structure. By analyzing local phase content, we proved that most structural information of the original natural scene is not lost after FEM-based whitening. By comparing results from FEM and flash-based acquisitions, we noticed that the active fixational strategy makes the event-based sensor to extract less redundant and still informative content. As a final beneficial consequence of whitening, reliability of phase information is expanded in a wider range of spatial frequencies^21^. We can hence conclude that fixational instability encourages redundancy minimization in neuromorphic vision by boosting the equalization of natural-images’ amplitude spectrum while preserving its phase spectrum and increasing its reliability at high spatial frequencies. In other words, FEMs contribute to an efficient encoding of the visual scene providing a pre-whitened signal to RGCs for further processing.

Finally, we analyzed the effects of possible biases in the direction of FEMs. Isotropy in the motion strategy was reflected on the acquired signal, leading to an equalization of sensor response to all oriented edges in the image. These results could possibly suggest an additional role of FEMs in biological systems - beyond those already postulated in the literature - related to their erratic nature, for which no theory has been advanced so far. Specifically, this equalization strategy could underlie an unbiased representation of image features carried out by orientation-selective cortical neurons, which is believed as one of the most important functionality of early vision, supporting subsequent object recognition and scene understanding.

For a long time, the idea of the eye operating as a standard camera (i.e. taking discrete spatial snapshots of the scene) has dominated visual neuroscience. However, alike other sensory modalities, vision is an active process in which the eye palpates external objects by means of motion. Movements transform spatial features into specific temporal modulations on the retina^26^, which consequently shape neural dynamic patterns in cortical regions. However, the results here presented concern on spatial information only, following a well-established framework for analyzing encoding efficiency (which mainly refers to the old “camera model” of the eye). To fully appreciate the functional role of FEMs, the organization of information in time should be addressed as well, since distribution of events in each sensor pixel is strongly structured by the motion sequence. Pure spatial information could be finely encoded in time as precisely synchronized activity of retinal neurons^27^, or phase-locked firing patterns across nearby cells^28^. Specifically, fine details of shapes, texture, and motion could be encoded by inter-cell temporal phases, instantaneous intra-burst rates, and inter-burst temporal frequencies of individual RGCs, respectively. Therefore, temporal dynamics provided by FEMs could similarly benefit neuromorphic vision applications. By productively spreading visual information in time, they could ultimately aid subsequent brain-inspired spike-based processing stages - able to learn complex temporal codes - to effectively extract rich informative content.

## Methods

### Set-up and data acquisition

The set-up for reproducing FEM-like motion on the event-based sensor is mainly composed of a remotely controlled motorized unit for the generation of precise pan and tilt rotations of the camera. Specifically, we use a *Pan-Tilt Unit* (PTU) E46 by FLIR Commercial Systems Inc. (as it provides smooth movements with a resolution as low as ∼0.8 arcmin) with a neuromorphic sensor DAVIS-346 (346 ×260 resolution, C-mount lens) from iniVation S.p.A. mounted on top of it (see Figure 6A). The DAVIS device provides both traditional gray-scale frames from an *Active-Pixel Sensor* (APS) and unconventional spiking events from a *Dynamic Vision Sensor*. Furthermore, this device also has a built-in *Inertial Measurement Unit* (IMU) with a (maximum) sampling frequency of 1 kHz and timestamps synchronized to the DVS events. An event is defined as a tuple **e**_*k*_ = {*t*_*k*_, **x**_*k*_, *p*_*k*_}where *t* represents the timestamp of the spiking event, **x**_*k*_ = (*x*_*k*_, *y*_*k*_) the location of the camera pixel sensing the event and *p*_*k*_ ∈{−1, +1} its ON/OFF polarity (*k* denotes the *k*^*th*^ spike). The mechanism for event generation can be summarized based on the *Brightness Constancy Equation*, which relates the temporal contrast (luminance derivative) to spatial contrast (image gradient) in the presence of relative sensor-scene movements^29^:

**Figure 6.**
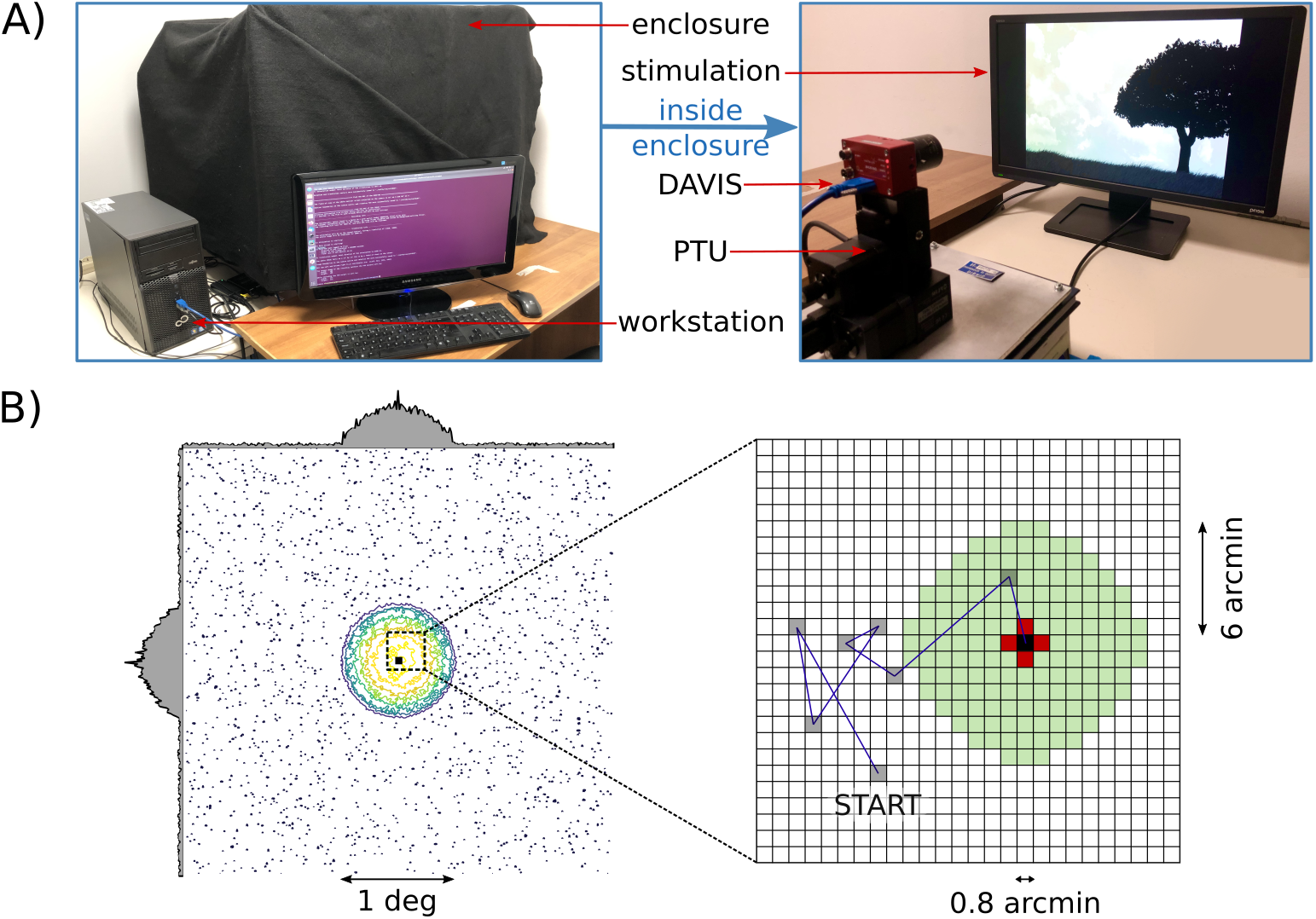
System setup for FEM-based visual acquisition. A) Left panel: the setup used for collecting data, as viewed from the user. We can identify the workstation for controlling all acquisitions, a utility display, and the enclosure in which the recordings are conducted. Right panel: inside view of the enclosure showing the DAVIS device, the PTU and the stimulation screen. B) Contour lines of a sample activation field from the adapted SAW model. The circular region of the foveola is delimited by lower activation values. The zoomed-up panel on the right comparatively illustrates the mechanism for deciding subsequent FEM steps of the original SAW model and our modified version. Gray-filled spots depict the history of a FEM sequence, with the final (current) lattice site shown in black. In the original model, the grid spot with the lowest activation value among the red-filled spots was chosen as the arrival site of the current step. In the adapted model, instead, the choice is among all the light-green lattice sites.

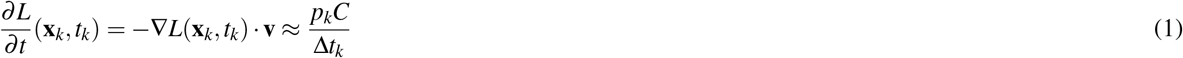

where *L* = log(*I*) is the log photocurrent (“brightness”), *C >* 0 is the temporal contrast threshold and Δ*t*_*k*_ is the time elapsed since the last event at the same pixel.

The PTU-DAVIS system was placed - at a fixed distance of 30 cm - in front of a BenQ LCD monitor with a 24 inches diagonal and resolution 1920 ×1080 @144 Hz. Such system was mounted on top of a mechanical platform having four rubber shock absorbers underneath for dampening the vibrations induced by PTU motors - which could otherwise propagate to the monitor causing its shaking. The whole setup was finally encapsulated inside a dark enclosure ensuring constant lightning conditions. Only the master computer (running Ubuntu 20.04 LTS) and a second display (used for experiment supervision) were left outside the enclosure. The described setup and all its components are shown in Figure 6A.

A custom-designed Python-based software pipeline was created to automatically conduct and efficiently control all acquisitions, providing a tool for finely tuning and significantly speeding up the data-collection procedure. The toolkit is therefore able to simultaneously deal with data transmission between a host computer and both peripheral devices, leveraging multiprocessing techniques. Specifically, it manages the communication with the PTU (for sending motion commands and receiving devices’ feedback), and with the DAVIS (for selecting bias parameters and logging the output-data in memory). Both communications were based on serial connection. Furthermore, either static images or synthetic visual stimuli (created with *PsychoPy*) could be reproduced on the monitor with controlled timings while recording data. Finally, the software also provides a tool for calibrating the position of the camera with respect to the stimulation display, thus finding the projection matrix for mapping 3D points in the world to 2D points in the camera plane. This was done based on the APS frames (given by the DAVIS) and functions from the *opencv* library. Finding the camera pose from known point correspondences on the monitor was useful for further processing data, mapping the ROI on the original image stimulus to the corresponding one on the camera plane.

### A model for FEMs

In order to simulate fixational eye movements for later driving the neuromorphic camera we used the *Self Avoiding Random-Walk* in a lattice model^12^. This model explicitly takes into account the fact that, in biology, the whole path of FEMs is confined to a small area: the foveolar region representing ∼ 1 deg of visual angle^30^. Movements are driven by a self-generated activation field and confined in foveola by a convex-shaped quadratic potential. The decision on the next step is based only on the sum between activation and potential of the four neighboring lattice sites with respect to the current one: the minimum is chosen and the activation at the current site is increased. As a result, the walker tries to avoid returning to recently visited sites and a self-organized distribution of activation over the lattice is built up. Therefore, the walker approximates *persistence* behavior of biological FEMs on a short timescale. After many time steps, the walker reaches lattice sites with high activation and potential values and is pushed back towards the center of the lattice, i.e. exhibiting *anti-persistence*.

It must be stressed, however, that the original model assumes the walker can only move along the two cardinal directions. We extended this range by defining a less constrained neighborhood (see illustration in Figure 6B): a circular window with radius equal to the maximum biological step size for a drift movement (i.e. ∼6 arcmin as reported by^31^, although precise measures are difficult to achieve^32^ and more recent studies suggest higher values^33^). The distance between adjacent points of the grid was set equal to the resolution of the PTU, since it is comparable to the minimum biological size of a drift step. The reason why we decided to relax the definition of the neighbourhood is that the walker (i.e. the sensor) can now explore all possible directions without being forced along the horizontal and vertical axes only. After we made these changes to the model, we checked whether persistence and anti-persistence behaviors were still discernible. We quantified them by estimating the *mean squared displacement* (MSD) of the random walk at a lag *l* between two iterations of the model:

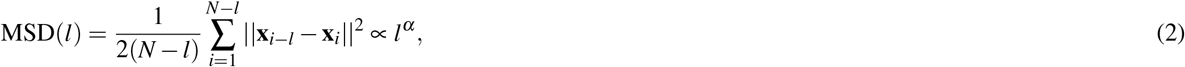

where *α* represents the power-law scaling exponent. In our analysis we set *N* = 10^4^ as the maximum number of FEM steps (iterations) of all the tested sequences and we averaged the results for 50 different motion seeds. Short and long timescales were defined as in^12^, i.e. *l* ≤ 10 and *l >* 10 iterations, respectively. Both the model and subsequent analysis were implemented in Python.

### Experiments

Three different sets of experiments have been conducted with the above described setup. The first one was required for the isotropy study, the second one for the amplification/whitening study, while the last one for all the other results. In all experiments, both DVS and IMU data have been recorded, with DVS biases at their default values. Pan/tilt speeds and accelerations have been set with the aim of achieving a fixed FEM-step frequency of ~ 50 Hz, as for the average frequency in the 0-100 Hz range of biological drifts and tremors^32^.

In the first set of experiments we used synthetic gratings as visual stimuli and recorded the sensor’s response with a single SAW-based motion pattern made of 40 steps. Two experiment sub-sets were performed based on the choice of gratings’ parameters: (i) 8 spatial frequencies evenly spaced between 0.2 and 1.6 cyc/deg with 12 orientations evenly spaced between 0 and 165° and maximum contrast value, and (ii) same as before but with contrast value changing with spatial frequency according to the 1*/k* statistics of natural images (where *k* represents spatial frequency), as in^9^. For these experiments, only events falling in the central 80*x*80 pixels region were recorded (all other events were filtered out from sensor’s FPGA). This was done for avoiding issues transmitting too many events at a time over the USB and is justified given the spatially homogeneous nature of grating stimuli (recording from larger regions is unnecessary because the stimulus does not change). Note that this region corresponds to 16 deg visual angle in both directions, ensuring that even the lowest spatial frequencies used were clearly visible (with enough cycles in such area).

For the second set of experiments we used natural images as visual stimuli, but with two distinct kinds of stimulation: (i) the sensor shaking in front of the monitor with SAW-based FEM sequences and (ii) the sensor staying still while stimulus flashed on the screen. In the former, natural images were kept on the screen for all the duration of the experiment. The program was then paused for ∼200 ms before the PTU started moving, and for a similar period afterwards. Six different random seeds were used for the FEM sequences and each of them consisted in 60 steps (∼1.2 s duration). During the latter experiment, instead, the monitor started displaying a gray screen. The gray value was set equal to the median intensity in the natural image used as stimulus, which appeared immediately on the screen after a ∼200 ms period. The image was then kept on the monitor for a maximum of 1.5 s or until data recording ended. This stimulation and recording procedure was repeated for six different trials. In both cases, the set of natural images used consisted of 16 samples taken from the van Hateren’s grayscale natural-image dataset^18^.

In the last set of experiments we adopted synthetic gratings again. However, this time we used both SAW-based (isotropic) and orientation-biased (anisotropic) FEMs, consisting in a total of 40 steps (∼0.8 s duration) in both cases. Anisotropic FEMs were modeled as random walks forced towards a specific orientation. The direction of each step was drawn from a peaked Gaussian distribution centered on the given orientation bias of the movement - defined on a [0, 180°) range - and a standard deviation of 10°. Step amplitudes, instead, were randomly selected from a uniform distribution in ±6 arcmin (negative values account for steps in [180°, 360°) range), with minimum step size equal to the PTU resolution as for the SAW-based model. A total of three directional biases were used for anisotropic FEMs (specifically, 30°, 60° and 90°) and four different seeds were tested for both types of movements. The set of biased movements used was chosen such that the resulting trajectories were confined in the same region as for SAW-based FEMs (since no explicit confinement of the whole sequence was defined in this case). Concerning the synthetic visual stimuli, we used a set of 12 differently-oriented gratings evenly spaced between 0 and 165°. Both spatial frequency and contrast were kept constant. Data from the sensor was only recorded from the central 80×80 pixels region, as in the first set of experiments.

### Events pre-processing

All pre-processings were handled by means of a custom Python repository. In all recordings of natural images, the 346 ×260 pixel matrix of the sensor was cropped to the central squared region of 200 ×200 pixels - since visual stimuli fell on a 40 ×40 degrees of sensor’s visual angle and outside this region data could be corrupted by the borders of the monitor. This region was smaller for recordings of synthetic stimuli, being 80 ×80 pixels, as previously mentioned. First, events were undistorted according to the camera matrix and distortion coefficients given by a previous camera calibration. Finally, hot pixels were identified and their events removed from all recordings.

For all the recordings with the moving sensor, we used IMU data to find the FEM time interval. This allowed us to only grub the events falling in that period. Specifically, angular speed was taken and smoothed with a Gaussian filter. The starting timestamp of the FEM interval was detected as the first sample where speed increased by four standard deviations from the mean value of the plateau (corresponding to the first ∼200 ms of recording, where the PTU had not been moving yet). Since all FEMs consisted of a fixed number of steps and frequency was set to 50 Hz, the average FEM duration was known. Therefore, such duration was considered for finding the last timestamp of the FEM interval in all recordings. Only events inside the detected interval underwent subsequent analysis. Some examples of recordings (both IMU angular speed and DVS instantaneous firing rate) are shown online in the Supplementary Fig.S2. The same figure displays how the DVS activity roughly reflects the FEM sequence followed by the sensor during acquisition: the firing activity is phase locked to the movement and its amplitude relates to that of the underlying FEM steps.

For static experiments, instead, the transition when the stimulus flashed on the monitor was detected in each recording by applying a similar analysis as above. However, this time the starting timestamp was found based on the average instantaneous firing rate across the whole pixel array (thus based on DVS information instead than IMU, since no movement was induced on the sensor in this case). As a matter of fact, the flash of the stimulus caused a sharp increase in the overall activity of the sensor, that gently decayed to the baseline after a while (∼ 200 ms).

### From events to frames

In order to compare event-based recordings of natural images with standard whitening procedures, we had to convert the event streams to analog signals (traditional frames). This can be achieved from a pixel-wise accumulation of event polarities over an arbitrary time interval Δ*t*. Doing so we produce an image Δ*L*(**x**) encoding the amount of brightness change that occurred during such interval^34^:

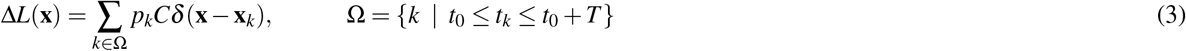

where *δ* is the Kronecker delta representing pixel **x**_*k*_ on the lattice and **x** is a generic pixel. All other terms are in accordance with equation 1. We adopt this conversion since it preserves knowledge on dark-to-light or light-to dark transitions, hence retaining spatial phase information from the event stream. In other words, ON (OFF) contrast polarities are encoded as positive (negative) values in the resulting image, while the amount of spatial contrast is encoded by the net number of spikes. Similarly, images achieved by applying traditional filter-based whitening techniques encode spatial contrasts polarity by the sign of the resulting convolution.

The appropriate time interval *T* in which to accumulate events can significantly change depending on the dataset, such that it is sometimes adapted to the amount of texture in the scene^34^. In order to find the optimal temporal window for our dataset of natural images we use mutual information^35^. Specifically, we tested 1000 progressively-increasing windows *T*, from a minimum of 10 ms and up to the whole FEM duration. We computed mutual information between the original image and the frame reconstructed from each time window, looking for the window *T* that maximizes it. The outcome of this process is summarized in the solid gray curve of Figure 7. As expected, mutual information is always increasing. This is reasonable since, while increasing the integration time, we incrementally add information to the event-reconstructed image. As argued in^36^, a short interval can lead to frames that are not sufficiently discriminative as they do not contain enough information, while an interval too long may wash out object contours due to motion. For this reason, the optimal time window could be found as the point where the average mutual-information curve visibly bends, namely the elbow or point of maximum curvature. This represents the best trade-off between the dimension of the integration window and the amount of information collected. Since the average curve was still noisy, we searched the elbow on its best fitting based on a truncated power law. This was done based on the *Kneedle* algorithm proposed in^37^, specifically by using the *KneeLocator()* functionality of the Python *kneed* package. The resulting optimal time interval was of 200 ms (see Figure 7). Based on the first 200 ms from the beginning of the stimulation, we therefore reconstructed a single frame for both FEM-based and flash-based experiments.

**Figure 7.**
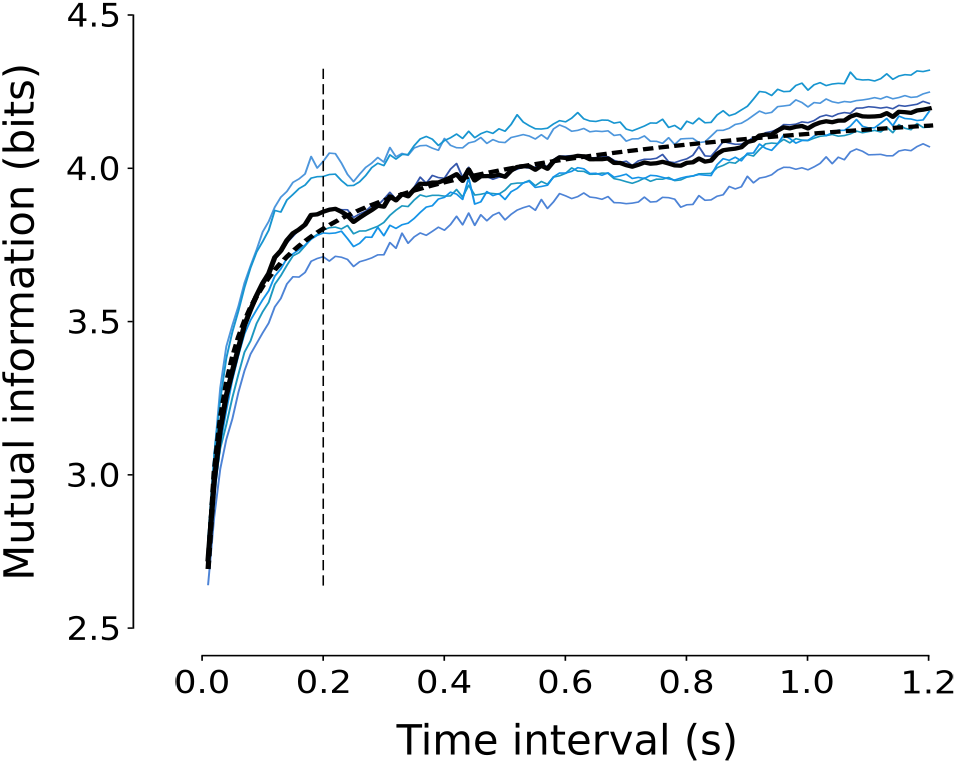
Trend of the mutual information between FEM-based reconstructed frames and corresponding natural images as a function of the time interval *T*. Blue lines show such trend for all six FEM seeds, the solid gray line the average across all seeds and the black dashed line its fitting. The dashed vertical black line represents the detected elbow point of the curve, i.e. the optimal time interval for frame generation (see text).

### Assessment of isotropy and spectral response

In order to characterize the overall system’s response to different orientations in the image for both biased and unbiased movements, we computed the mean firing rate of all sensor pixels as a function of stimulus orientation, by averaging over all spatial frequencies and movement seeds. Similarly, the overall spectral characterization of the system was computed as the firing rate of sensors pixels for each spatial frequency tested and by averaging over all orientations.

Concerning the isotropy of motion sequences, we quantified it based on the circular statistics of the Euclidean distances - relative to the whole path length - travelled in each of the 12 directions considered for stimuli, up to 180°. The computation was limited on the [0, 180°) range since isotropy had to be related to the effect that each FEM step caused on the perception of a specific oriented contrast, independently of contrast’s direction. In other words, a movement along an arbitrary orientation *θ* has the same effect on the net firing activity of the system as a movement (with same length) towards *θ* + 180°, if events’ polarity is disregarded. In our experiments, we tested four different sets of movements (i.e. four populations: three biased plus one unbiased), each one having 240 samples (i.e. four seeds per 60 steps). A Rayleigh test^38^ was then computed on their weighted distributions of angles (remapped on the full circle). Furthermore, we also performed the symmetry test around the median and, since in our case the mean direction of biased movements was known in advance, the V-test for circular uniformity^39^ (the median direction was used for the unbiased population). All statistical tests were performed based on the Python implementation of the *“CircStat”* MATLAB toolbox^40^, *“pycircstat”*.

### Decorrelation strategies

Due to the 1*/k*^*n*^ falloff of natural images’ amplitude spectrum (with *k* the radial spatial frequency, and typically 1 ≤*n* ≤1.3), the variance along the low-frequency eigenvectors is much larger than the variance along the high-frequency eigenvectors. This produces huge differences in the variance along different directions. Some techniques have been proposed for ameliorating these effects and thus decorrelating natural image signals. A well-known method is inspired by the receptive field properties of retinal ganglion cells^41^. In such a model, the image is convolved with a simple spatial linear filter: a *Difference of Gaussians* (or DOG), which corresponds to a band-pass filter in the frequency domain. This filter is composed of the difference of two radially-symmetric Gaussian kernels sharing the same center but having different standard deviations. For our analysis, we set the standard deviations of the inner and outer Gaussian kernels as 0.7 and 1.12 pixels, respectively. Based on these parameters we chose 13 pixels as the optimal spatial support of the filter.

We also considered a second method, based on the whitening procedure proposed in^16^ (which we refer to as OFW in the text). Here, a cascade of two filters is applied to the Fourier spectrum of the image: a whitening (high-pass) filter and a smoothing (low-pass) filter. The whitened image is then obtained through inverse Fourier transform. The goal of the whitening filter is to explicitly counterbalance the statistics of natural images, thus its frequency response is *W* (*k*) = *k*^*n*^. In order to design the appropriate filter for matching the statistics of our natural images, a linear regression model was applied on the profile of images’ amplitude spectrum in logarithmic scale (azimuthal average across eight directions and averaged across the whole set). The resulting amplitude profile was best fit by *n* = 1.15. The implementation of such a filter alone yielded a roughly flat amplitude spectrum across all spatial frequencies. However, since high frequencies are typically corrupted by noise and aliasing, it is not wise to boost them indiscriminately. For this reason, the low-pass filter with frequency response *L*(*k*) = exp(*k/k*_0_)^4^ was used (with a cutoff frequency *k*_0_ of 2 cyc/deg). On the whole, the cascade of these filters resemble the spatial-frequency response characteristics of retinal ganglion cells (i.e. the DOG model).

Before applying such whitening techniques on the original natural image, we cropped and down-sampled it in order to match the size and shape of event-based recordings (i.e. 200 *×* 200 pixels). The auto-correlations were then computed for all images based on 400 random squared patches with side of 6 deg (i.e. 30 *×* 30).

### Phase-based analysis

For this analysis we used the very same images as in the previous case but focusing on FEM- and flash-based recordings. As a reference signal for phase information we use the OFW-filtered image. In order to compute local phase, we designed a bank of Gabor filters with six orientations (*θ*_*q*_, *q* = 1 … 6) equally spaced between 0 and 165° and 29 spatial frequencies equally spaced in the range [0.2, 2) cyc/deg, with one octave relative bandwidth. By combining the responses from basis channels with different orientations, but a common frequency, we derived information about local energy, local phase and dominant local orientation around each pixel location (**x** = (*x, y*)) of the image, according to the formulation proposed in^42^. Specifically, the local energy associated to each orientation at a given spatial frequency was computed as:

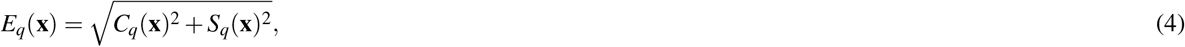

where *C*_*q*_ and *S*_*q*_ are the image convolutions with even and odd components (respectively) of a complex Gabor filter oriented along *θ*_*q*_. The *dominant local orientation* was derived as:

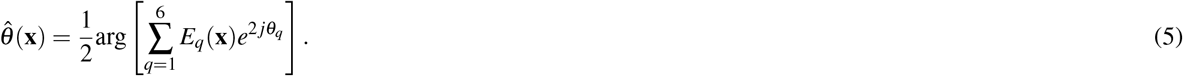

The multichannel even and odd Gabor filter responses were obtained by interpolating the single orientation channel responses in the dominant orientation:

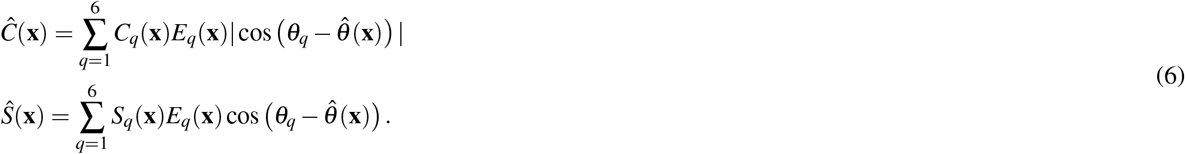

From these components, we straightforwardly derived the *dominant local phase* 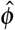 as:

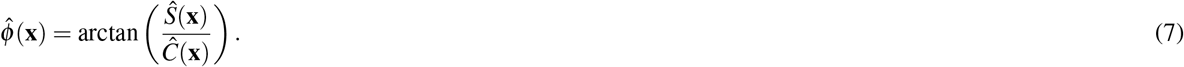

Finally, we had to pick up a metric for comparing the dominant local phase of event-based frames with that of a reference whitened frame at all the given scales of the Gabor bank. To this aim, the *Phase Locking Value* (or PLV) was chosen. PLV is commonly used in EEG studies for evaluating the synchronization between two signals^20^. Here, we use it as a metric for analyzing the consistency of the detected dominant local phase in a given signal with respect to a reference. This decision is justified by the invariance property of such metric to some constant phase shift over the entire pixel array. Conversely, a phase-similarity metric would give us low values if local phase information differs in the two images by some constant value, even though the phase structure is basically preserved. As a matter of fact, a global phase shift could be present in the data gathered from the neuromorphic sensor, possibly due to a small misalignment of the event-based frames and the original natural stimulus. However, the presence of a constant phase shift does not invalidate the preservation of images’ phase structure in the events stream: it does not reflect some alteration of the original structure and therefore must be tolerated. For these reasons, the PLV measure - formally defined below - seems as the most appropriate to our situation:

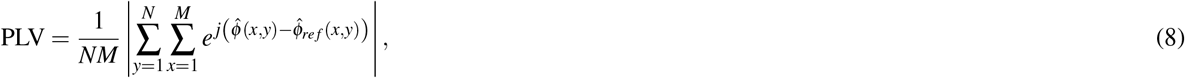

Where 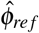 represents the reference dominant local phase - given by the OFW-based image (reference signal) - and 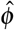 is related to the event-based acquisition process in exam. As discussed above, a unitary PLV means local phase of the recording procedure is consistent with that in the OFW method. A zero PLV value means there is no local-phase congruency. Anyhow, the resulting PLV should be compared to that of a set of surrogate images where phase structure has been randomly altered. Specifically, for a given pair of OFW-filtered image and FEM-based (or flash-based) reconstructed image, a set of 100 surrogates was built by randomly selecting an (*x, y*) location on the latter frame and swapping the four resulting image patches with respect to such coordinate. Doing so, for each image pair and each scale (spatial frequency), we obtain a PLV of the unaltered image reconstructed from events and a distribution of the PLV from 100 different versions of the same image not preserving the phase content. Finally, at all filter scales, we computed the average (and standard deviation) PLV across all 16 visual stimuli and six FEM seeds (or flash trials). For FEM-based and flash-based surrogates, instead, we compute the 95^*TH*^-percentile across all visual stimuli, FEM seeds (or flash trials) and all 100 samples.

### Equivalent filters of step movements

Our aim was to characterize the effect of single FEM steps on the visual information acquired by the sensor. FEM patterns are composed of many independent steps from which a single frame can be isolated in the resulting event stream. We therefore divided all FEM-based recordings of natural images in 60 non-overlapping time windows of 20 ms, roughly equivalent to the duration of a single FEM step (given the FEM-step frequency imposed to the PTU). For each time window we built the corresponding frame according to the procedure presented in the events pre-processing section above. Since each step originates slightly different sensor responses, isolating different characteristics of the underlying image, we can assimilate each of them to a specific convolutional linear operator acting on the original image *s* and producing its filtered version *o*:

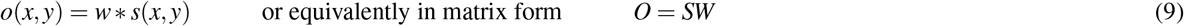

where *W* is a column vector with *n*^2^ elements defining the weights of the *n×n* convolution kernel, *S* is a circulant matrix where each row represents the flattened version of a single *n×n* patch of the original image *s, O* is the flattened column-vector version of the output image *o*. The idea is to calculate the convolution kernel based on the original image and its convolved version. The best approach is to build it as an optimization problem in the spatial domain where we want to find the weights of the convolving kernel *w* minimizing the sum of squared residuals. Due to the possible ill-posed nature of the problem, we opted to include a regularization term in the minimization, favouring a solution with smaller norm:

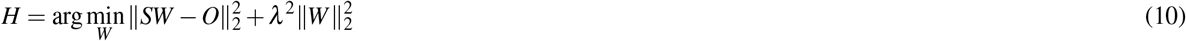

This sort of minimization problem is the standard form of Tikhonov regularization, or ridge regression, where λ is the regularization parameter that balances the influence of the first term (fidelity) and the second term (regularization). Using the regularization term allows us to control also the smoothness of the weights. The above Tikhonov minimization problem has a unique solution given by:

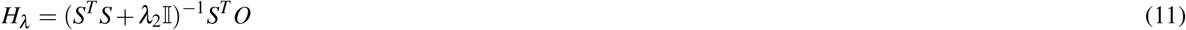

where 𝕀 is the identity matrix. By applying equation 11 to each of the 60 frames reconstructed from the event-based recordings, we can estimate the equivalent filter *H*_*λ*_ associated to each single step.

We considered 3000 random patches of 13 ×13 pixels (∼2 deg visual angle) from each of the 60 FEM-based reconstructed images and averaged the resulting filters across all 16 natural stimuli. The regularization parameter *λ* was set to 20. The size of the patch was chosen according to the spatial support of the DOG filter defined above. Hence, we achieve a set of 60 filters for each FEM seed. Note that frames were previously standardized, i.e. subtracting their mean and dividing by their standard deviation. By computing the average filter across all 60 results, we obtain an equivalent filter relative to the whole FEM sequence. Finally, a Fourier representation with 100 ×100 pixels (zero-padding size of 87 pixels in both directions) was built from all such filters for visualizing their effect in the frequency domain.

## Data availability

The data generated and used in this study will be made available from the corresponding author upon request.

## Acknowledgements

Research reported in this publication was partially supported by the National Eye Institute of the National Institutes of Health under Award Number R01EY032162. The content is solely the responsibility of the authors and does not necessarily represent the official views of the National Institutes of Health.

## Author information

### Contributions

All authors conceived the work together with all experiments. S.T. and A.C. acquired and analyzed data, while S.P.S. assisted in the interpretation of results. The first draft of the manuscript was written by S.T., who revised it together with A.C. and S.P.S. Financial support for the project was provided by S.P.S.

