## Supplemental Figure 1 for "Active Fixation as an Efficient Coding Strategy for Neuromorphic Vision"

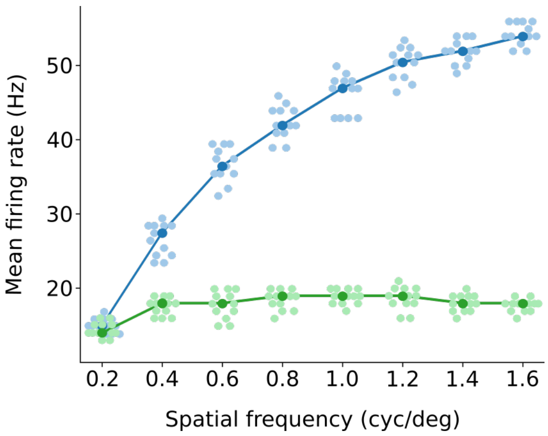


**Figure 1.** Comparison of the mean firing rate evoked from sinusoidal synthetic stimuli at different spatial frequencies in a range $[0.2, 1.6]$ cyc/deg in a condition where contrast is kept constant across frequency (blue line) or it is varied according to a power-law behavior of natural images (green line). Dark color dots represent the average across the 12 tested orientations of grating stimuli, single data points are displayed in light colors. We applied an horizontal jitter to single data points for the sake of visualization. Blue curve shows an amplification effect from the system, while the green one shows an equalization (whitening) of its response to varying spatial frequencies. This whitening effect is attributable to the opposing trend of natural image distribution across spatial frequencies with respect to the amplification introduced by FEMs. An amplification of system's firing activity at increasing stimulus' spatial frequency is visible when grating's contrast is kept constant (blue curve). On the contrary, a whitening effect (response equalization with spatial frequency) is shown if contrast is adjusted with spatial frequency according to natural image statistics (green).
