## Supplemental Figure 2 for "Active Fixation as an Efficient Coding Strategy for Neuromorphic Vision"

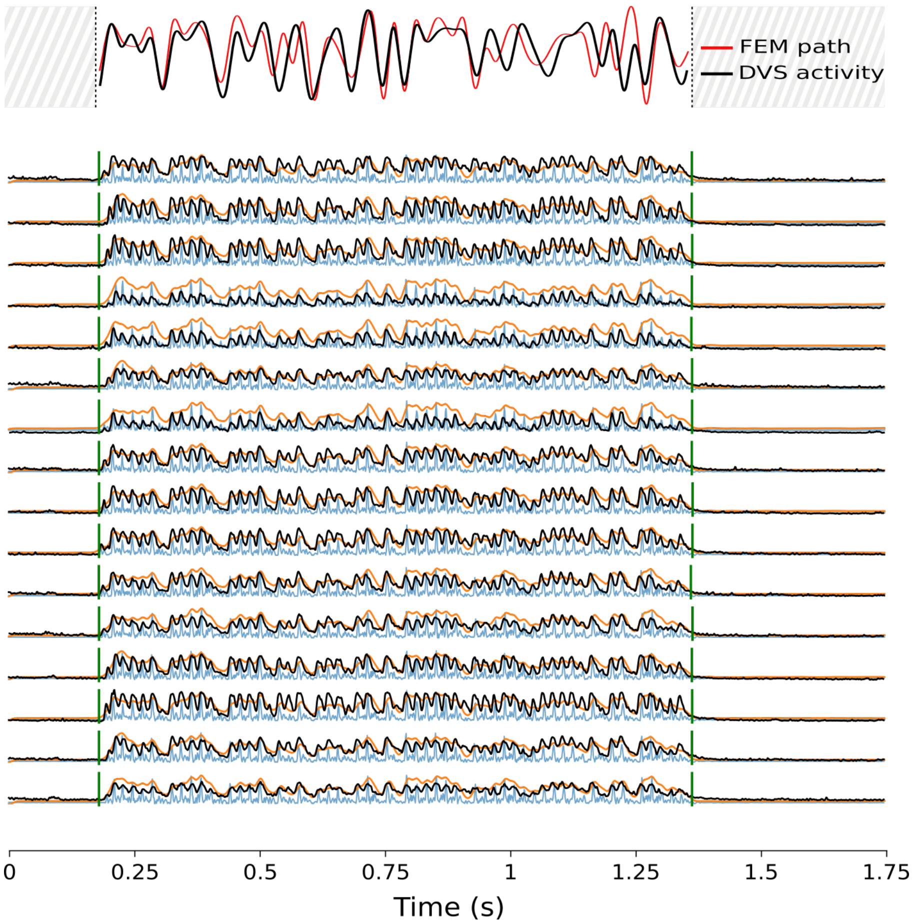


**Figure 1.** Examples of FEM sensor activity and IMU angular speed recordings. (Top) the distance covered by the walker during a specific FEM sequence (specific seed) is shown in red (interpolation over 60 FEM steps); the (smoothed) instantaneous firing rate of the DVS (averaged across 16 different recordings) is superimposed in black, sampled at 60 steps and interpolated. (Bottom) The 16 plots show both IMU angular speed (raw signal in light blue, smoothed signal in orange) and DVS firing activity (black). All recordings are relative to the same FEM sequence but different visual stimuli. All signals have 1 kHz sampling frequency.
